# Chemogenetic activation of parvalbumin interneurons in the piriform cortex alleviates hippocampal focal seizures

**DOI:** 10.1101/2023.12.22.572953

**Authors:** Huiqi Zhang, Zhuo Huang, Chunyue Geoffrey Lau

**Affiliations:** Department of Neuroscience, College of Biomedicine, City University of Hong Kong, Hong Kong SAR, China; Shenzhen Research Institute, City University of Hong Kong, Shenzhen, China; State Key Laboratory of Natural and Biomimetic Drugs, Department of Molecular and Cellular Pharmacology, School of Pharmaceutical Sciences, Peking University Health Science Center; Beijing, 100191, China

**Keywords:** Temporal lobe epilepsy, Long-range modulation, Seizure therapeutics, Piriform cortex, Seizure network

## Abstract

**Background:** Temporal lobe epilepsy (TLE) involves aberrant changes in the brain at molecular, cellular and circuitry levels, leading to recurrent seizures. Approximately one-third of patients develop drug resistance. Circuit-based interventions show promise for alleviating the drug-resistant seizures. This study explored the efficacy of chemogenetic stimulation on specific neuronal populations in piriform cortical microcircuits in a mouse TLE model.

**Objective:** The anterior piriform cortex (APC) is a limbic area closely associated with TLE but is understudied. Our previous study demonstrated that in the APC, parvalbumin-expressing (PV^+^) interneurons provide strong inhibition and help maintain excitation-inhibition balance. Here, we examined whether and how APC^PV^ neurons can ameliorate seizure symptoms in TLE.

**Methods:** We used a chronic intrahippocampal kainic acid (IHKA) mouse model that develops hippocampal pathology and spontaneous recurrent seizures (SRSs). APC^PV^ neurons were chemogenetically activated, and local field potentials were recorded longitudinally from multiple regions. Seizure metrics, spectral power and functional connectivity were compared between APC^PV^-activated and control samples.

**Results:** Loss of PV^+^ synapses emerged in the APC during epileptogenesis. Selective activation of APC^PV^ neurons reduced the frequency and duration of hippocampal SRSs and modified long-range dynamics, with region- and frequency-specific changes in interictal band power and functional connectivity.

**Conclusion:** APC^PV^ chemogenetic activation exerts a robust anti-seizure effect and identifies the APC as a promising cortical node for seizure network control. The associated spectral and connectivity signatures suggest that targeted modulation of APC microcircuits can rebalance distributed seizure networks and lower seizure recurrence.

## Introduction

Epilepsy is a debilitating neurological disorder exhibiting recurrent seizures, affecting 8 per 1000 population lifetime prevalence, with 61 per 100,000 new cases added every year [1]. Among its various forms, temporal lobe epilepsy (TLE) is the most common type of focal epilepsy. While TLE can be treated with antiepileptic drugs, one third remain resistant to medication [2]. Surgical resection of epileptic foci, though effective in controlling seizures, carries significant risks and major side effects [3]. In contrast, neurostimulation-based approaches offer key advantages, such as non-destructive intervention, reversibility, and adjustable modulation. Accumulating evidence suggests that TLE not only results in pathological changes in the foci, usually the hippocampus and surrounding structures, but also in remote structures [4]. Understanding the roles of these interconnected regions and specific cell types is therefore essential for both deciphering TLE pathophysiology and developing targeted neuromodulation therapies.

The anterior piriform cortex (APC), a key component of the olfactory system and limbic circuitry, has emerged as a critical player in TLE. While extensively studied for its role in odor processing, the APC possesses unique anatomical and physiological properties that make it particularly susceptible to seizure activity [5–7]. The region is characterized by strong recurrent excitatory connections among principal neurons (PNs), which normally support associative memory functions but can lead to pathological hyperexcitation under epileptic conditions [8,9]. Moreover, the APC is heavily connected with the nearby endopiriform nucleus, also termed the “area tempestas”, which has exhibited the lowest threshold for kindling seizures [10,11]. Experimentally, large lesions of the piriform cortex (PC) can retard seizure acquisition in the rat kindling model [12]. Clinically, surgical resection of the PC in TLE patients has been associated with improved seizure outcomes [13]. Additionally, vagal nerve stimulation, an effective neurotherapeutic, may exert its benefits by modulating the synapses of PC [14]. These studies cement the notion that the PC is an important player in seizure generation and TLE treatment.

At the cellular level, the APC, much like the hippocampus, contains PNs and a diverse population of GABAergic interneurons that maintain excitatory-inhibitory (E-I) balance [15,16]. Cell-specific chemogenetic methods have been pivotal in epilepsy research to investigate the mechanisms and to identify cell populations as therapeutic targets [17]. Most chemogenetic interventions improve the E-I balance in epilepsy by (1) activating interneurons using ligand-binding engineered G-protein coupled receptors, hM3D(Gq) or (2) silencing PNs with hM4D(Gi)- or KORD(Gi)- chemogenetic tools [18]. From a therapeutic development perspective, we propose targeting neuronal populations whose activation (rather than inhibition) can suppress seizures - an approach particularly relevant for clinical translation to electrical stimulation therapies. Our prior work demonstrated that within APC microcircuits, PV^+^ interneurons (but not SST^+^) exert potent inhibition capable of gating cortical output [19]. However, the seizure-suppressing potential of APC^PV^ interneurons remains to be systematically investigated in epilepsy models.

In the present study, we employed an established mouse model of TLE induced by intrahippocampal kainic acid (IHKA) injection, which reliably produces chronic epilepsy with focal spontaneous recurrent seizures (SRSs) originating from the sclerotic hippocampus [20]. Using a combination of chemogenetic approaches and multi-site electrophysiological recordings, we investigated the effects of selectively activating APC^PV^ interneurons on SRSs. Our results demonstrate that APC^PV^ chemogenetic activation significantly reduces the frequency and duration of SRSs while inducing widespread changes in interictal network dynamics. These findings established a crucial anti-seizure role for APC^PV^ interneurons and provided important insights into potential network mechanisms underlying their therapeutic effects.

## Materials and methods

### Mice

All experimental procedures were approved by the Animal Research Ethics Sub-Committee of City University of Hong Kong and conducted under a license granted by the Hong Kong SAR Government Department of Health. PV-Cre mice (Jackson Laboratory, stock #008069) of both sexes were used in this study. To minimize confounding variables, experimental groups were balanced for body weight, age, and sex. Mice were group-housed in the Laboratory Animal Research Unit (LARU) at City University of Hong Kong under a 12-hour light/dark cycle (lights off: 20:00–08:00) with ad libitum access to food and water.

### Intrahippocampal Kainic acid (IHKA) chronic model

Six- to seven-week-old mice were anesthetized with a ketamine/xylazine (KX) mixture (100 mg/kg and 10 mg/kg, respectively). KA (0.5 µg/µL, 500 nL; Alomone Labs, Cat #K-200) was stereotaxically injected into the right dorsal hippocampus (dHPCR) at the following coordinates (relative to bregma): AP: -2.2 mm, ML: 1.8 mm, DV: -2.1 mm. To minimize liquid runaway, the glass pipette was left in place for 10 minutes post-injection. Mice typically recovered from anesthesia within 60 minutes. Status epilepticus (SE) was confirmed by observing mild convulsions, drooling, immobility, or unidirectional rotational movements. SE self-terminated within 2–8 hours. Post-injection, mice were checked daily for seven days to assess overall health.

### Immunohistochemistry and image acquisition

Brain sections were prepared for immunofluorescent staining of PV protein (rabbit anti-PV, 1:1000, Swant, PV27a) or c-Fos protein (rabbit anti-c-Fos, 1:500, Abcam, ab190289), electrode location verification and virus expression validation. Images for virus and electrode location verification were acquired using an upright fluorescent microscope (Nikon Eclipse Ni-E). High-resolution images for PV^+^ synapses were acquired using confocal microscopy (Nikon A1HD25 high speed and large field of view confocal microscope). Quantitative analysis of PV^+^ synaptic puncta was conducted using FIJI/ImageJ software. Complete protocols for immunohistochemistry and image analysis are provided in the Supplementary Methods.

### Virus injections and electrode implantations

Electrodes were implanted into multiple regions for local field potential (LFP) recording. Mice underwent both virus injection and electrode implantation five weeks after IHKA-induced SE. Mice were deeply anesthetized with KX mixture. Tissue paper was put under the animal’s body to prevent heat loss during the surgery. The head was shaved and fixed in a stereotaxic instrument (Stoelting, 51730). After incision and skull exposure, the skull surface was wiped clean with sterile PBS. Using stereotaxic coordinates relative to bregma, we injected 400 nl of either AAV-hSyn-DIO-hM3D(Gq)-mCherry (Addgene #44361-AAV5) or AAV-hSyn-DIO-mCherry (Addgene #50459-AAV5) (titer 7×10¹² vg/mL) bilaterally into the anterior piriform cortex (APC; AP: +1.54, ML: ±2.6, DV: -4.5 mm). Virus was delivered at 2 nl/sec through a glass pipette (1.5 µl, Drummond Wiretrol) pulled using a vertical puller (Narishige PC-100), with the pipette remaining in place for at least 10 minutes post-injection to minimize runaway.

We then implanted custom-made single electrodes targeting right APC (APCR; AP: +1.5, ML: +2.6, DV: -4.5 mm), right dorsal hippocampus (dHPCR; AP: -2.2, ML: +1.8, DV: -2.0 mm), and bilateral primary somatosensory cortex (S1; AP: -1.2, ML: ±3.0, DV: -0.5 mm). Electrodes were assembled with Teflon-coated stainless-steel wires (114.3 µm coated diameter, A-M Systems, 790600) inserted into silica tubing (inner diameter: 150 µm, outer diameter: 220 µm, Fisher Scientific, 0624461). Two stainless steel screws were placed on the skull above cerebellum to serve as reference and ground electrodes. All electrodes and screws were connected to a six-pin micro-connector, which was then secured to the skull with dental cement.

Post-operative care included subcutaneous administration of carprofen (10 mg/kg) for three days to manage pain and inflammation. We allowed three weeks for full viral expression before conducting experiments. Electrode positions and viral expression were verified through post-hoc histological examination.

### Longitudinal LFP recording of chronic IHKA model

LFPs were recorded from freely moving mice during the chronic epileptic phase, 56–65 days after IHKA injection. We conducted a 9-day recording protocol with chemogenetic manipulation during the middle three days. Animals received intraperitoneal injections of either saline or Clozapine N-Oxide (CNO; 3 mg/kg, HelloBio HB6149) to selectively activate APC^PV^ neurons. Recording sessions began 30 minutes post-injection to allow for optimal drug absorption and neuronal activation. We acquired signals using a Medusa recording system (Bio-Signals Technologies) that simultaneously captured 4 channels of LFP at 1000 Hz sampling frequency with 16-bit digital resolution. To account for potential circadian influences on seizure expression, recording schedules were strictly controlled. Each mouse underwent Pre-CNO, CNO, and Post-CNO recordings at a consistent time of day relative to the 12-hour light/dark cycle. Day sessions began at 09:00, and night sessions began at 20:00. Throughout the recording period, mice had unrestricted access to food and water while freely moving in the recording chamber. This naturalistic setup allowed us to monitor seizure activity under conditions that closely resembled the animals’ normal living state.

### Focal seizure quantification

LFP data were processed using custom MATLAB scripts. Signals were baseline-corrected and bandpass filtered (1-200 Hz) using MATLAB’s fir1 function. Epileptic spikes were detected when the negative peak prominence exceeded 3 standard deviations of the signal (using MATLAB’s findpeaks function). Focal seizures (FS) were defined as spike clusters meeting two criteria: (1) spike frequency >2 Hz and (2) duration >10 s. This FS identification is based on previously established criteria [21]. To validate the robustness, we conducted a sensitivity analysis by varying the detection parameters. Specifically, we tested negative peak prominence thresholds ranging from 2.5 to 3.5 SD, FS durations from 5 to 15 s, and spike-rate criteria from 1 to 3 Hz. Across all tested permutations, the longitudinal pattern of detected FS remained consistent. The stability of these results across varying detection stringencies confirms that our findings are not dependent on specific algorithmic parameters. Then, we quantified three parameters: FS number, total seizure time, and individual FS durations. For statistical comparison of FS durations between conditions, we acquired the estimated median differences in Pre vs CNO, and Post vs CNO, with 95% confidence intervals using 5000 bootstrap samples. Generalized seizures are not stably expressed. If it shows, the subsequent 30 min of LFP recordings were manually excluded from analysis because such events cause a prolonged suppression of neuronal activity.

### Analysis of interictal spectral power and functional connectivity

We extracted 60 random 10-second interictal trials per animal per day from the LFP recordings, excluding any segments containing amplitudes exceeding 500 µV. Each trial comprised 4-channel signals that were prepared to compute both spectral power and phase-based connectivity measures.

The power spectrum is the magnitude squared of Fourier transform of data (*fft* function in Matlab). We acquired the trial-averaged power spectrum for each mouse. The absolute band power was the area under the power spectrum curve in the corresponding frequency range (*trapz* function in Matlab). We examined six canonical frequency bands: δ (1–4 Hz), θ (4–8 Hz), α (8–13 Hz), β (13–30 Hz), low-γ (30–50 Hz), and high-γ (50–90 Hz). Power analysis was conducted for each animal and each brain region.

The phase locking value (PLV) was used to quantify the strength of neural synchronization, also called functional connectivity. PLV can be calculated according to this formula: 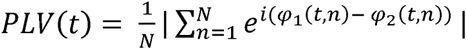, where N is the number of trials and □_1_ and □_2_ are the instantaneous phase of two channel signals at time t and trial n. The *pn_eegPLV* function was used in this analysis (https://www.mathworks.com/matlabcentral/fileexchange/31600-phase-locking-value). Two LFP signals were first filtered in a specific frequency band, then computed with Hilbert transform to obtain the instantaneous phase value, □. The trial-averaged PLV was calculated for each mouse. After that, a ten-second segment of instantaneous PLVs was averaged along the time axis. Therefore, we obtained a single PLV measure for each region pair per frequency per animal. PLVs from the same region pair, frequency band and condition (Pre/CNO/Post) were averaged across animals (see analysis pipeline in Fig. 3). These averaged PLVs were visualized in three-layer connectivity graphs. The arithmetic difference of PLV (ΔPLV) was obtained by subtracting one layer from another (CNO - Pre, or Post - CNO).

### Principal component analysis (PCA) and support vector machine (SVM)

We employed principal component analysis (PCA) to reduce dimensionality and visualize patterns in the LFP features. The dataset comprised 45 samples (5 animals × 9 days), with each sample characterized by 60 interictal features including: regional spectral power (4 regions × 6 frequency bands) and inter-regional phase locking values (6 region pairs × 6 bands; see Supplementary Table 1 for complete feature list). After standardizing the data, PCA identified the most informative feature combinations. To assess condition separability while preserving independence at the subject level, we trained a support vector machine (SVM, scikit-learn package in Python) using the first three principal components as inputs and performed leave-animal-out cross-validation. In each fold, data from one animal were held out as the test set and the model was trained on data from the remaining animals. Model performance was quantified as the mean classification accuracy for discriminating CNO versus non-CNO conditions across folds.

### Statistics

Statistical analyses were performed in MATLAB or Python with appropriate methods as indicated in results and figure legends and in a double-blind manner. Comparisons of three parametric data sets were performed either with a one-way ANOVA or with repeated measures (RM) ANOVA, in the case of matched groups. If an ANOVA indicated that there exists significant difference among groups, Tukey’s multiple comparisons test was performed additionally. When the assumption of normality for the repeated measures design was violated, a non-parametric Friedman test was utilized to compare measurements across the three groups. Significant main effects were followed up with Wilcoxon signed-rank tests with Bonferroni correction for pairwise comparisons. Unless otherwise specified, data were shown as mean ± sd. We did not use statistical methods to predetermine sample size. No outliers or other data were excluded. For all analysis, a p-value < 0.05 was considered statistically significant. The significance is indicated as * p < 0.05, ** p < 0.01, *** p < 0.001 and **** p < 0.0001. The details of statistical tests and results were shown in the results and figure legends. All figures were prepared using MATLAB, Python and Illustrator CC (Adobe).

## Results

### PV^+^ synapses were weakened in the HPC and APC in the chronic IHKA model

How does inhibition change in the epileptic brain? We examined PV^+^ synapses in the chronic IHKA model. Four weeks after KA injection, brains were fixed and PV immunostaining was assessed in the APC and two hippocampal (HPC) subregions, the dentate gyrus (DG) and CA1. KA administration markedly reduced PV stained area in all regions (Fig. 1). In the HPC, the ipsilateral side showed the strongest loss: perisomatic PV^+^ synapses in the DG granule cell layer (GCL) and CA1 stratum pyramidale (SP) were largely absent, and the contralateral side also exhibited a substantial reduction (Fig. 1A,B,D). In the APC, PV^+^ synapses in layer 2 (L2) declined similarly on both sides, whereas PV^+^ somata were preserved and remained concentrated in L3 (Fig. 1C,D). These changes support the idea that inhibitory synapses are highly vulnerable during epileptogenesis. [22].

**Figure 1.**
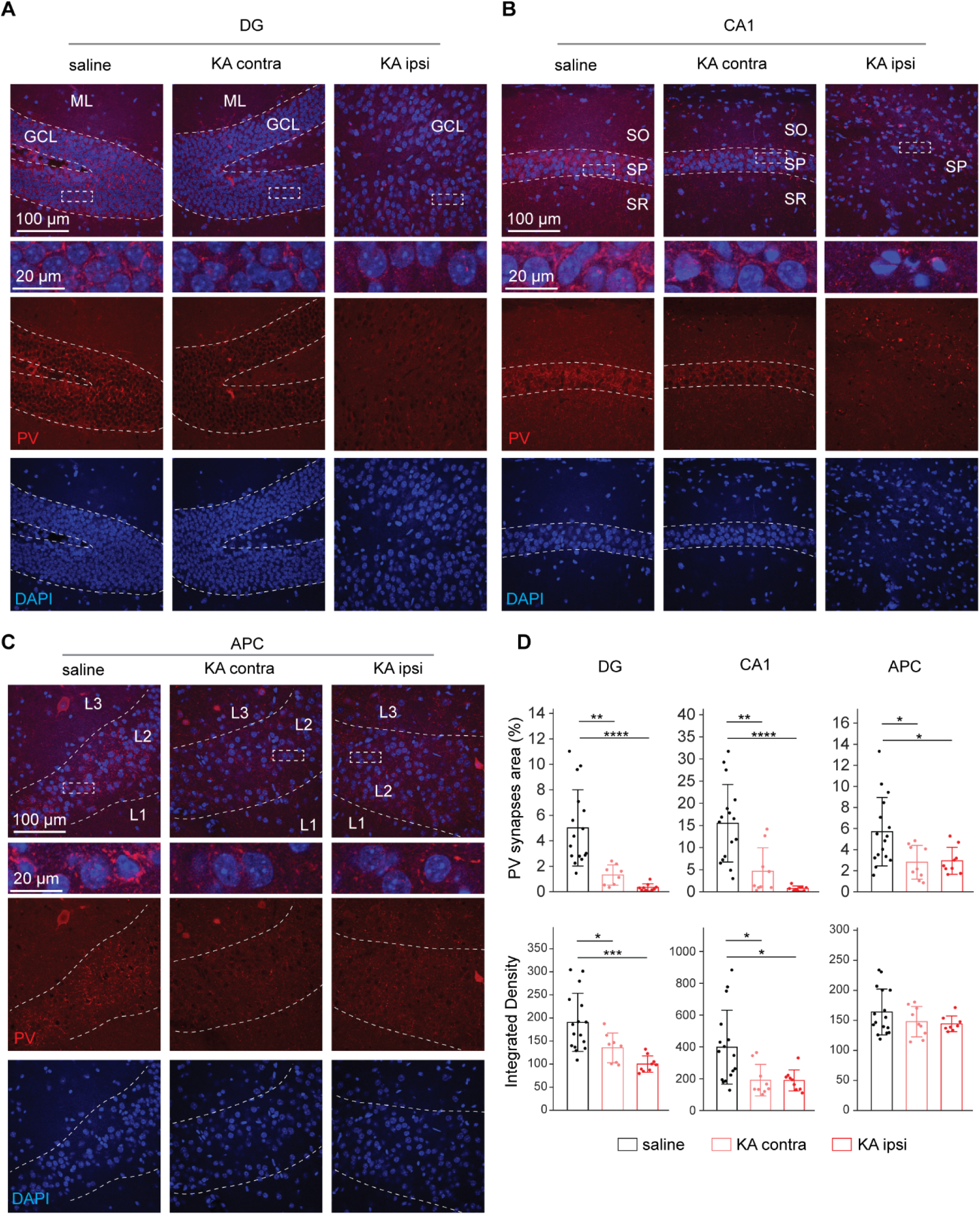
PV^+^ synapses were reduced bilaterally in the HPC and APC in the chronic IHKA model. (**A**) Representative images of PV^+^ synaptic puncta in the dentate gyrus (DG) granule cell layer (GCL) under three conditions: saline injection, contralateral (contra), and ipsilateral (ipsi) sides relative to KA injection. (**B**) Representative PV^+^ expression in the CA1 stratum pyramidale (SP). (**C**) Representative PV^+^ expression in APC layer 2 (L2). (**D**) Quantification of PV synaptic expression. Top: the percentage of PV^+^ synapses covered area in DG (saline: 5.01 ± 2.99, n = 16; KA contra: 1.31 ± 0.79, n = 8; KA ipsi: 0.35 ± 0.28, n = 9), CA1(saline: 15.49 ± 8.76, n = 16; KA contra: 4.65 ± 5.30, n = 9; KA ipsi: 0.79 ± 0.55, n = 9) and APC (saline: 5.70 ± 3.25, n = 16; KA contra: 2.80 ± 1.60, n = 9; KA ipsi: 2.94 ± 1.28, n = 9). Bottom: integrated density of PV^+^ synapses in DG (saline: 190.46 ± 62.83, n = 16; KA contra: 135.25 ± 32.19, n = 8 ; KA ipsi: 100.01 ± 17.78, n = 9), CA1 (saline: 398.14 ± 232.47, n = 16; KA contra: 190.70 ± 98.69, n = 9; KA ipsi: 188.82 ± 66.01, n = 9) and APC (saline: 164.02 ± 38.16, n = 16; KA contra: 147.83 ± 25.19, n = 9; KA ipsi: 144.28 ± 13.16, n = 9). One-way ANOVA with Tukey multiple comparison as post hoc, * p < 0.05, ** p < 0.01, *** p < 0.001, **** p < 0.0001. Data were shown as mean ± sd. n indicates the number of sections. N = 3 mice for each condition.

Consistent with previous work, principal neurons in DG GCL and CA1 SP became dispersed and lost their tightly packed organization after IHKA injection [23], with a similar dispersion of neurons in APC L2. To assess potential neurodegeneration, we quantified cell density. In DG and CA1, DAPI^+^ cell density decreased only on the ipsilateral side, whereas in the APC it was significantly reduced bilaterally compared with saline controls (Supplementary Fig. 1). Thus, in the chronic epileptic brain, synaptic loss and neurodegeneration extend beyond the hippocampal focus. In the APC, PV^+^ neurons were largely preserved but PV^+^ synapses were downregulated, implying weakened local inhibition and a lowered threshold for SRSs. We therefore hypothesized that reactivating APC PV neurons could counteract this vicious cycle and reduce seizure occurrence.

### Selective activation of APC^PV^ neurons attenuates seizures in chronic IHKA model

We used chemogenetic stimulation based on Designer Receptors Exclusively Activated by Designer Drugs (DREADDs) to selectively activate PV^+^ neurons in the APC. An excitatory DREADD, hM3Dq, was expressed in APC^PV^ interneurons by injecting a Cre dependent AAV into PV Cre mice. Clozapine N oxide (CNO) was administered to activate hM3Dq. Immunofluorescent staining of PV protein in APC sections showed that 99.5% (n = 596 cells) of hM3Dq-mCherry-expressing neurons were PV^+^, indicating highly specific DREADD expression, and 94.0% of PV+ neurons expressed hM3Dq, demonstrating efficient transduction (Supplementary Fig. 2A). Co labeling of c Fos and hM3Dq confirmed robust activation of infected neurons by CNO (Supplementary Fig. 2B).

We next combined chemogenetic stimulation with electrophysiology to test how APC^PV^ activation affects SRSs (Fig. 2A). In the chronic phase (5 weeks after KA), hM3Dq was delivered bilaterally to the APC, and electrodes were implanted in the APCR, dHPCR, and bilateral S1 for local field potential (LFP) recordings (Fig. 2B); hM3Dq expression and electrode tracks in the APC are shown in Fig. 2C. After three weeks of viral expression, animals underwent nine recording days, receiving CNO from days 4–6 and saline on the remaining days. Stable focal seizures (FSs), defined as spike clusters lasting >10 s at >2 Hz, were observed in dHPCR (Fig. 2D,E), with epileptiform activity present in all four channels; the dHPCR electrode track is shown in Fig. 2F. In PV hM3Dq mice, CNO significantly reduced FS number and total FS time, which returned to baseline after CNO withdrawal (Fig. 2G,H), and also shortened individual FS duration (Fig. 2I). In PV mCherry control mice, CNO had no effect on FS number or duration (Fig. 2J–L), indicating that the anti seizure effects in PV hM3Dq mice are DREADD dependent rather than off target CNO actions. Together, these findings show that activating APC^PV^ neurons decreases seizure recurrence and mitigates seizure severity.

**Figure 2.**
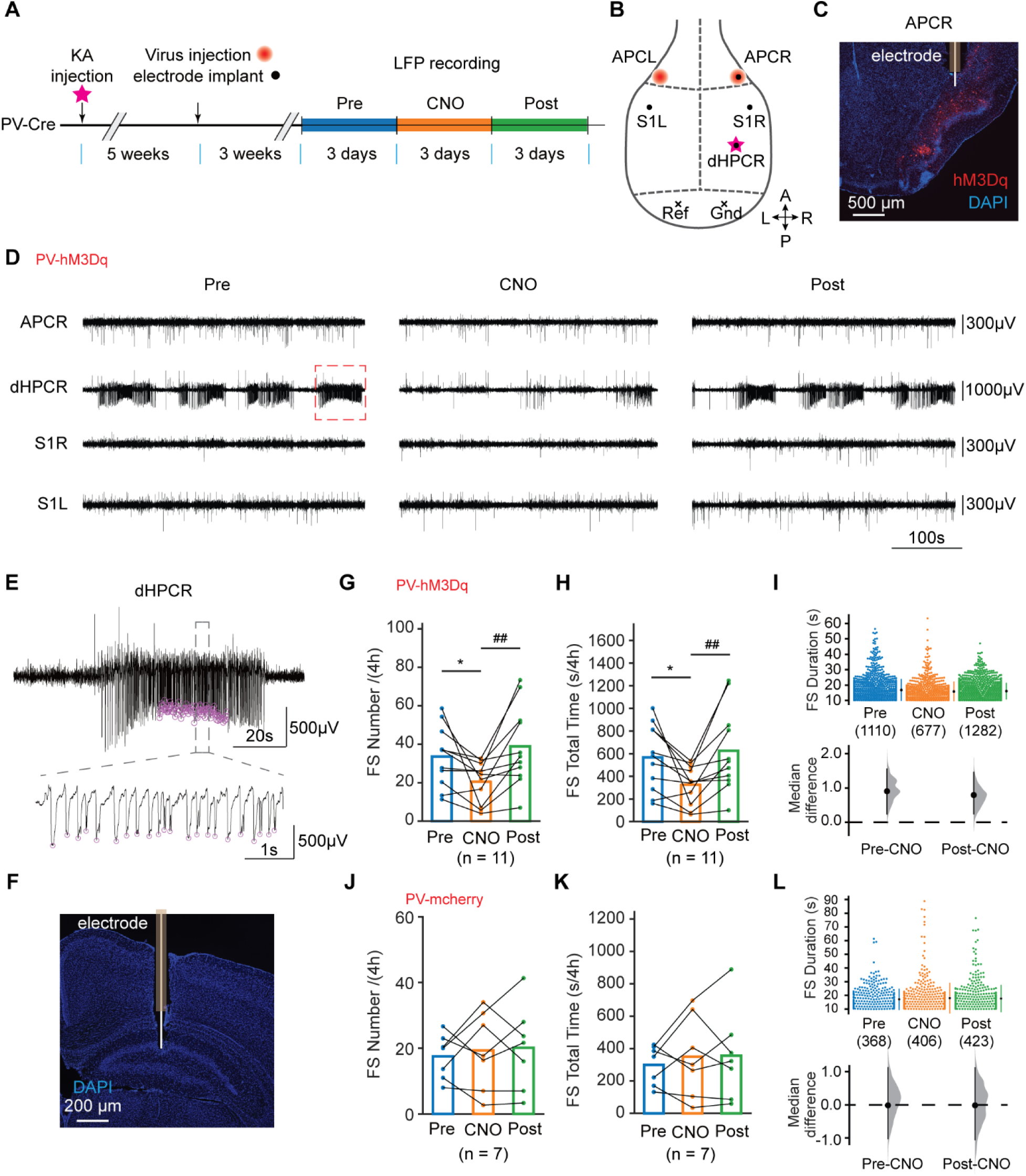
Chemogenetic activation of APC^PV^ neurons attenuated epileptic seizures in the chronic IHKA model. (**A**) Schematic of LFP recording and CNO treatment in KA-induced epilepsy with spontaneous recurrent seizures (SRSs). Virus (DIO-hM3Dq-mCherry or DIO-mCherry) was injected in bilateral APC and electrodes were implanted 3 weeks before longitudinal recordings. (**B**) Surgery schematic: KA injection site (magenta pentagram), bilateral APC virus injections (red circles), recording electrodes in APCR, dHPCR, S1R and S1L (black dots) and reference and ground locations marked (black crosses). (**C**) Representative DIO-hM3Dq-mCherry expression in APC^PV^ neurons with electrode placement. (**D**) Representative LFP traces of the four recording channels. From dHPCR channel, chemogenetic activation of APC^PV^ suppressed focal seizures (FSs). (**E**) Expanded view (red box in D) showing that FS defined as ≥2 Hz epileptic spiking (magenta circles) and lasting ≥10 sec. Bottom: High-temporal resolution trace of FS. (**F**) Representative image of the electrode track in the dHPC. (**G, H**) In the PV-hM3Dq group (n = 11 mice), chemogenetic activation of APC^PV^ neurons significantly reduced the number (G) and total time (H) of FSs in the chronic IHKA mice. Friedman test was used. Post-hoc analyses were subsequently conducted using Wilcoxon signed-rank tests with Bonferroni correction applied to adjust for multiple comparisons. *p < 0.05, compared between Pre and CNO groups. ## p < 0.01, compared between CNO and Post groups. (**I**) Chemogenetic activation of APC^PV^ neurons reduced the individual FS durations. Top: Swarmplot of individual seizure durations across Pre/CNO/Post conditions. Bottom: Bootstrapped median differences (circle) with 95% confidence intervals (vertical lines). (**J,K,L**) PV-mCherry controls (n=7): CNO had no effect on FS frequency (J), total time (K) or FS durations (L). Statistical test methods were the same as in G, H, I correspondingly.

### APC^PV^ neuronal activation alters long-range interictal network dynamics

Our results so far indicated that activation of APC^PV^ neurons exerts robust anti-seizure effects. Epileptic seizures are complex phenomenon whose dynamics depend strongly on large-scale network architecture [24,25]. Therefore, we further quantified power in each region and the functional connectivity between all region pairs to investigate how APC^PV^ activation affects the seizure network. We focused on interictal phase as it is relatively long and present a crucial window to prevent seizure recurrence. First, we extracted four-channel interictal segments from the LFP recordings (Fig. 3A). Functional connectivity was quantified using the phase locking value (PLV), which measures the consistency of phase relationships between signals over time. [26]. The analysis pipelines for power and PLV were implemented separately (Fig. 3B,C and Fig. 3D–H). Intuitively, if signals in two channels rise and fall together or maintain a stable phase offset across epochs, the PLV is high, indicating strong synchronization.

**Figure 3.**
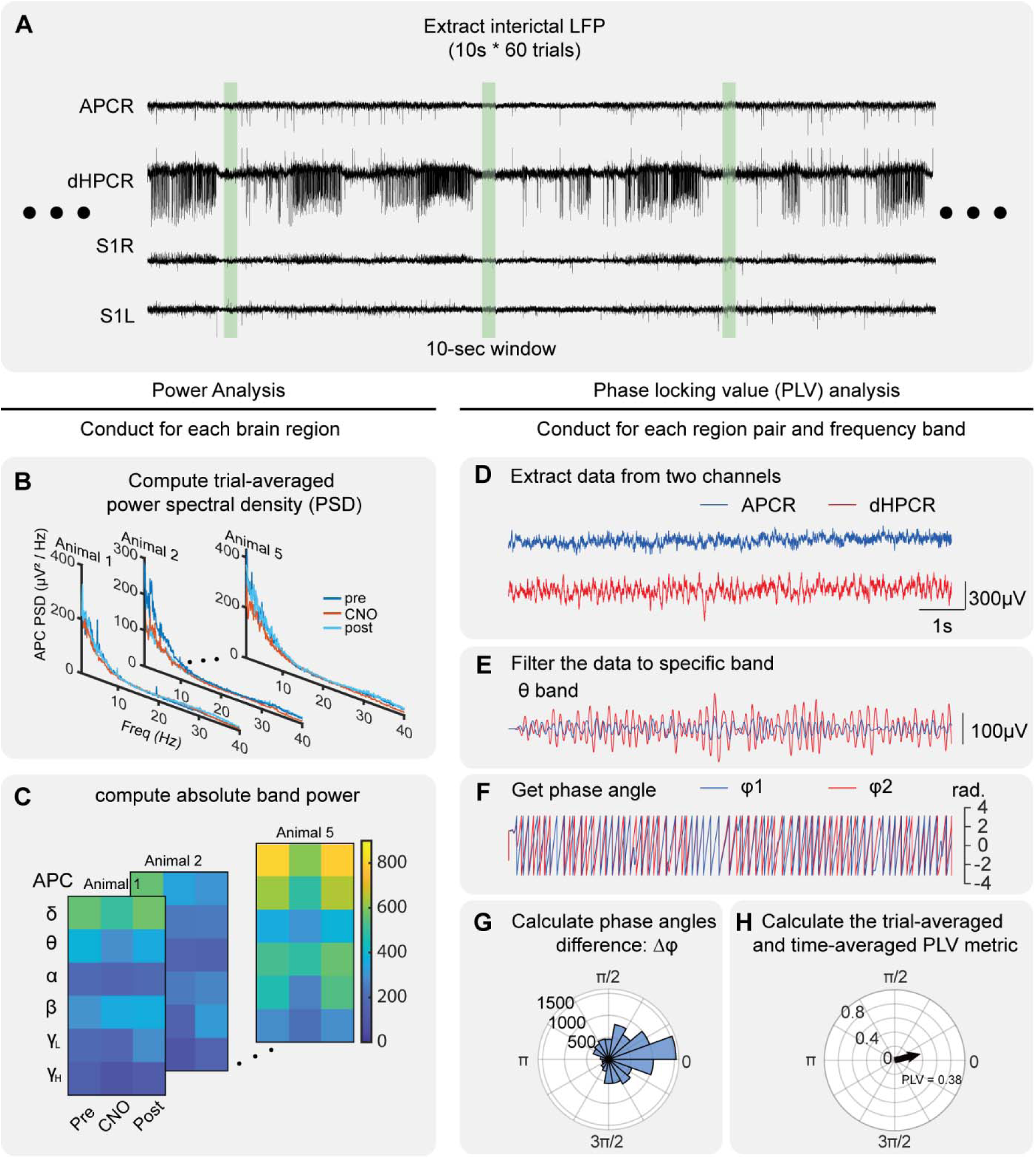
Network dynamics analysis pipeline during interictal periods. Spectrum power analysis pipeline is shown in panels B – C and connectivity analysis pipeline is shown in panels D – H. (**A**) Signal extraction: For each recording session (per animal per day), 60 interictal 10-second trials were extracted. (**B**) Power spectral density (PSD) was computed and averaged across trials for each animal and condition. (**C**) Absolute spectral power was calculated for six frequency bands for each condition, displayed as a heatmap (rows: frequency bands; columns: Pre, CNO, and Post conditions). (**D–H**) Functional connectivity analysis: (**D**) Representative raw LFP traces from two channels (shown: APCR and dHPCR). (**E**) Bandpass filtering of signals into specific frequency bands. (**F**) Instantaneous phase were acquired for each channel. (**G**) The histogram of phase angle differences in one trial. (**H**) Trial-averaged and time-averaged PLV was calculated for each region pair, frequency band and treatment conditions.

Interictal power changes were region- and band-specific (Fig. 4B). High-frequency power decreased in the APCR and HPCR when APC^PV^ neurons were activated, whereas low-frequency power (δ and θ) increased in bilateral S1. For functional connectivity, CNO treatment in APC^PV-hM3Dq^ mice differentially modulated PLVs across region pairs and frequency bands (Fig. 4C–E): it enhanced PLVs in APC connected pairs and/or high-frequency bands (e.g., APCR–dHPCR θ, APCR–HPCR high γ, APCR–S1 high γ), but reduced dHPC–S1L and S1L–S1R connectivity in low-frequency bands, indicating desynchronization in the δ, θ and α ranges. In PV mCherry controls, CNO produced no significant changes in either band power or connectivity (Supplementary Fig. 3). Thus, APC^PV^ chemogenetic activation induces heterogeneous, structured changes in interictal power and coupling.

**Figure 4.**
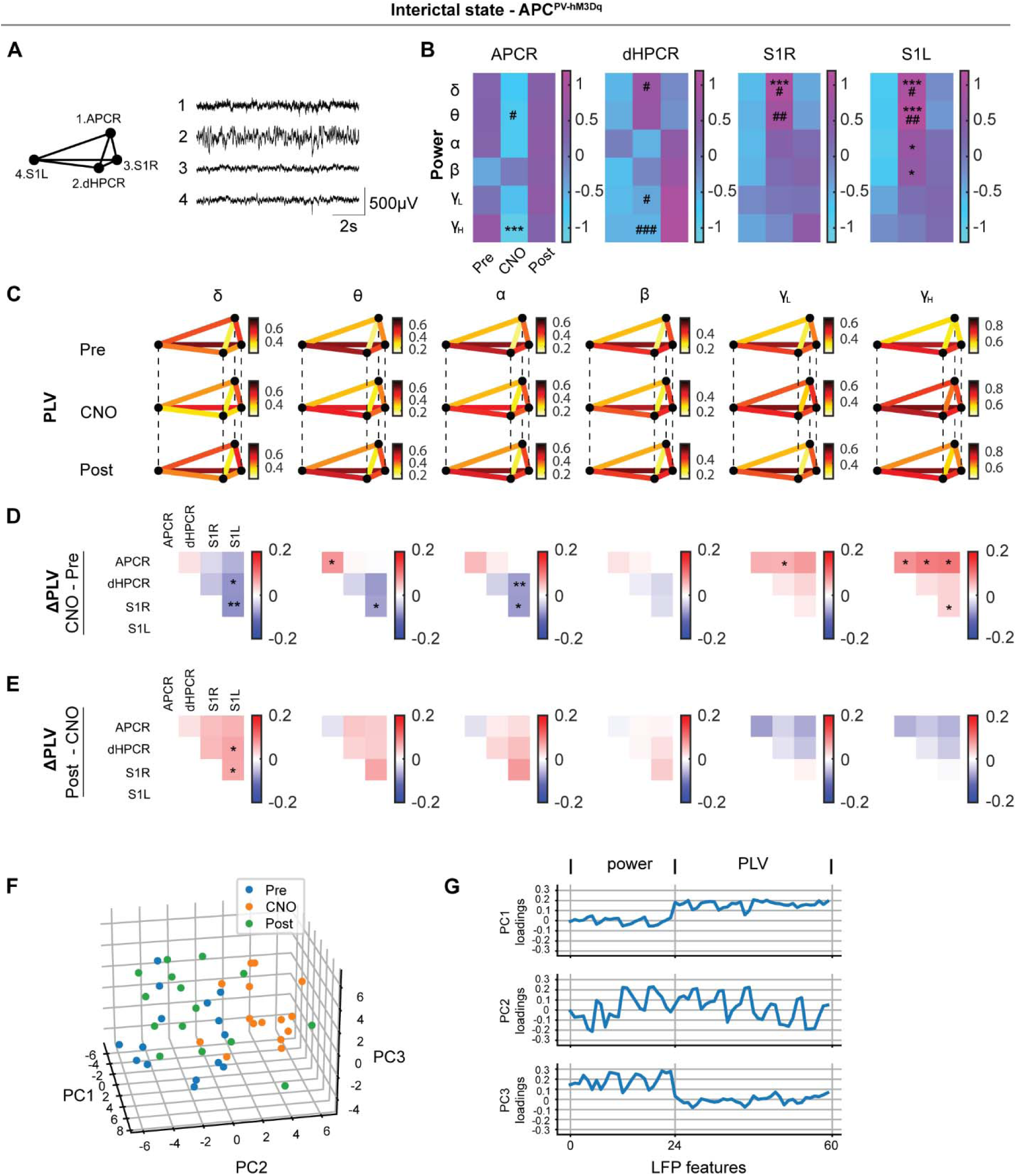
APC^PV^ neuronal activation alters long-range network dynamics during interictal periods. (**A**) Representative interictal LFP traces from four recording sites. (**B**) Normalized band power (z-scored within subjects for each channel and frequency band) during interictal periods displayed as a heatmap (rows: frequency bands; columns: Pre, CNO, and Post conditions; colors: mean band power; n = 5 mice). One-way repeated measures ANOVA followed by Tukey’s multiple comparison as post hoc test. *p < 0.05, ***p < 0.001, compared CNO with Pre. ^#^p < 0.05, ^##^p < 0.01, ^###^p < 0.001, compared CNO with Post. (**C**) Phase locking values (PLVs) between all region pairs across 6 frequency bands and Pre, CNO, and Post conditions (mean values, n = 5 mice). (**D, E**) ΔPLV matrices showing functional connectivity changes between: Pre and CNO conditions (D), CNO and Post conditions (E). One-way repeated measures ANOVA followed by Tukey’s multiple comparison test. * p < 0.05, ** p < 0.01. (**F**) Principal component analysis (PCA) of LFP spectral power and connectivity features. Each point represents a recording session (per animal per day) in PC space (Pre: blue; CNO: orange; Post: green), showing that CNO-treated samples were distinguishable from non-treated Pre and Post samples. (**G**) Loading weights displayed the contribution of original LFP features (detailed in Supplementary Table 1) to each principal component.

These findings implied that the “normal looking” interictal EEG contain information about the underlying network’s response to the therapeutic stimulation. We next asked whether these patterns are sufficiently systematic to distinguish treatment from non treatment days. To test this, we pooled all 60 interictal LFP features (regional band power and PLVs) and applied principal component analysis (PCA), treating each animal-day as one sample. The first three principal components (PCs) captured 62.3% of the total variance (PC1, 29.5%; PC2, 20.5%; PC3, 12.3%). In this low-dimensional space, CNO samples were clearly separated from Pre and Post samples along PC2, whereas Pre and Post overlapped. A support vector machine with leave one animal out cross validation achieved 0.844 accuracy in classifying CNO versus Pre/Post combined (Fig. 4F). PC2 reflected mixed contributions from power and PLV features (Fig. 4G). Together with the reduction in FS number (Fig. 2G), these findings indicate that APC^PV^ activation likely exerts seizure-preventing effects by jointly modulating local neural activity and inter regional synchronization.

## Discussion

The present study identifies APC^PV^ interneurons as a functional node of seizure control in a chronic mouse model of TLE and characterizes the network-level consequences of their chemogenetic activation. We report three interrelated findings: (1) PV^+^ synapses are lost in the APC as well as in the hippocampus during epileptogenesis in the chronic IHKA model; (2) bilateral chemogenetic activation of APC^PV^ neurons with hM3Dq reduces the frequency and duration of hippocampal SRSs in a DREADD-dependent manner; and (3) this targeted cortical intervention reorganizes interictal spectral power and functional connectivity across multiple brain regions in a structured, regionally heterogeneous pattern. From the interictal LFP features, treatment status, CNO versus non-CNO days, can be classified. Together, these findings extend seizure network theory by establishing that an extrahippocampal cortical population is capable of suppressing hippocampus-driven SRSs and that the interictal state encodes information about cortical inhibitory tone that is modulated by circuit-level stimulation.

The bilateral depletion of PV^+^ perisomatic synapses in APC L2 demonstrated an underrecognized vulnerability of fast-spiking interneurons beyond the canonical hippocampal focus [7,27]. Meanwhile, we observed that PV^+^ somata in APC L3 were preserved. This implies that inhibitory deficit in the APC reflects synaptic downregulation rather than frank neurodegeneration. This is a therapeutically important distinction, because cells that are intact but functionally silenced are amenable to reactivation, whereas neurons that died are not. The bilateral reduction in the APC stands in contrast to the predominantly ipsilateral hippocampal changes, raising the possibility that spread of epileptiform activity drives a progressive weakening of APC inhibitory circuits in the absence of direct KA injection [28]. This vicious cycle - whereupon hippocampal SRSs deplete APC inhibitory synapses, further reducing cortical gating of limbic re-excitation - motivated our hypothesis that restoring APC^PV^ function could interrupt seizure recurrence.

The robust reduction in SRS frequency, total seizure time, and individual seizure duration following APC^PV^ chemogenetic activation constitutes the central finding of this study. These effects were strictly DREADD-dependent: CNO administered to mCherry control animals produced no change in any seizure metric, excluding off-target pharmacological actions of CNO as a confound [29]. Targeting PV interneurons with excitatory DREADDs in the hippocampus has previously been shown to attenuate TLE seizures [21]. What distinguishes the present findings is that therapeutic activation was at a cortical site, the APC, whose PV interneurons are not immediately within the hippocampal seizure initiation zone. The efficacy of this extrahippocampal intervention demonstrates that seizure networks in TLE are not confined to the hippocampus and its immediate structural neighbors, but extend to cortical microcircuits capable of regulation of limbic excitability [30]. From a translational standpoint, targeting a cortical node such as the APC rather than the hippocampus itself may reduce the risk of cognitive side effects - particularly the memory impairment frequently associated with direct hippocampal interventions in mesial TLE patients [3].

The mechanistic basis for APC-mediated seizure suppression is likely to involve gating of excitatory output from the APC to the entorhinal cortex and, through multisynaptic pathways, to the dentate gyrus. In the APC, PV-expressing interneurons form strong perisomatic inhibition onto L2 PNs, and their output largely accounts for recurrent inhibition that controls APC cortical output, as our prior work demonstrated [19]; perisomatic targeting is uniquely positioned to control somatic spike generation and thereby gate cortical output [31,32]. When hM3Dq is activated, increased firing of these PV cells constrains the activity of L2 PNs, reducing the excitatory drive dispatched from the APC along the piriform-entorhinal projection toward the hippocampus. Consistent with this view, recent work has shown that disruption of the piriform-entorhinal-dentate trisynaptic circuit, whether by targeted circuit silencing or by pharmacological blockade, is sufficient to abolish kindling-induced generalized seizures [33]. The reversibility of these effects upon drug withdrawal confirms that the anti-seizure benefit was due to active neuronal engagement rather than long-lasting synaptic plasticity associated with DREADD expression itself.

APC^PV^ activation reshaped the interictal network in a manner that was both spatially heterogeneous and frequency-specific, providing mechanistic insight into how distant circuit modulation propagates through the seizure network. The reduction in high-frequency power in the APCR and dHPCR reflects decreased local excitability in both the modulated cortical node and the principal site of seizure generation, consistent with the view that high-gamma activity in these regions is a marker of network hyperexcitability and proximity to seizure threshold [34,35]. The increase in low-frequency power in bilateral S1 may represent propagation of the dampening effect to secondary cortices, or alternatively a change in the globally correlated slow-wave state driven by reduced hippocampal sharp-wave and ripple output. The connectivity analyses revealed that APC^PV^ activation enhanced phase synchrony between the APC and other regions in high-frequency bands, suggesting that stronger inhibitory control reshaped, rather than isolated, its network interactions. From a clinical neuroscience perspective, these observations align with the growing body of evidence that interictal functional connectivity encodes the epileptogenic state and its susceptibility to therapy [36–38]. The finding that multivariate interictal LFP features could distinguish CNO treatment days from baseline with 0.844 accuracy without reference to any seizure event implies that beneficial changes in network state are detectable in the interictal EEG. Such interictal biomarkers derived from multi-epoch spectral and connectivity analysis offer a potential avenue for tracking therapeutic efficacy across sessions and guiding parameter adjustments in adaptive neurostimulation strategies.

The present findings have direct implications for circuit-based neuromodulation strategies in drug-resistant TLE. Preclinical work has shown that low frequency electrical stimulation of the APC in KA treated rats markedly reduces spontaneous seizure frequency and abolishes generalized seizures [39]. It would be interesting to test whether APC^PV^ neurons are involved in the low-frequency stimulation benefits. Manipulation of olfactory cortex principal neurons bidirectionally influences seizure dynamics [40]. Our results suggest that the PV interneuron population of the APC is a key cellular effector through which piriform circuit stimulation exerts its therapeutic benefit, and that approaches designed to preferentially recruit inhibitory interneurons - including high-frequency stimulation paradigms or spatially targeted deep brain stimulation - could be more efficacious than non-selective excitatory approaches [41]. Besides electric stimulation, chemogenetic suppression of seizures has also been demonstrated in rodent and non-human primate models of seizures [42,43], further supporting the translational potential of this circuit-based strategy. The APC’s positioning as a major afferent gateway to limbic structures makes it an attractive substrate for responsive neurostimulation systems.

Several limitations of the present study warrant acknowledgment. While mCherry controls confirm DREADD specificity, some degree of CNO back-metabolism to clozapine cannot be fully excluded at the systemic doses used; adoption of next-generation agonists such as compound 21, which are more potent and have reduced off-target pharmacology, would further strengthen causal inference in follow-up work [29]. The IHKA model reliably creates hippocampal sclerosis and chronic SRSs but may not fully recapitulate the network diversity of human mesial TLE, in which sclerosis pattern, dual pathology, and network reorganization can vary substantially across patients. Our four-channel recording configuration does not capture intermediate nodes such as the amygdala, entorhinal cortex, or contralateral hippocampus that may participate in the observed connectivity changes. Further work is warranted to decipher the precise circuit mechanisms and provide a more complete picture of network reorganization.

In conclusion, this study establishes APC^PV^ interneurons as both a critical component of the TLE seizure network and a viable therapeutic target whose activation suppresses hippocampal SRSs and restructures interictal brain dynamics. These findings position the APC as a promising cortical node for seizure network control and suggest that circuit-based stimulation strategies targeting inhibitory interneurons in this region merit systematic clinical investigation.

## Acknowledgement

We would like to thank the Hong Kong Research Grants Council for their funds (RGC/ECS 21103818 to C.G.L., and RGC/GRF 11104320 and 11104521 to C.G.L.), internal funds from City University of Hong Kong (9610354 to C.G.L.), and National Natural Science Foundation of China funds (82271498 and 82341246 to Z.H.).

## Author contributions

H.Z. performed experiments, analyzed data and wrote the first draft of the manuscript. H.Z., Z.H. and C.G.L. reviewed and revised the manuscript. C.G.L. conceived and supervised the study and obtained funding. All authors read and approved the final manuscript.

## Data availability

The data and custom code that support the findings of this study are available from the corresponding author upon reasonable request.

## Supplementary material

Supplementary material is available.

