## Supplemental tables and figures for "Chemogenetic activation of parvalbumin interneurons in the piriform cortex alleviates hippocampal focal seizures"

### Supplementary method

##### Immunohistochemistry

Mice were deeply anesthetized and perfused with PBS and 4% paraformaldehyde. The whole brain was post-fixed overnight in 4% PFA followed by cryoprotection in 30% sucrose in PBS for 72 hours at 4°C. Brain sections (35 μm) were prepared with cryostat (HM525 NX, Thermo Fisher Scientific) for immunofluorescent staining or electrode location verification and virus expression validation. For immunohistochemistry, the sections were first washed in PBS three times for five minutes each time, blocked with 10% goat serum and 0.3% Triton x-100 in PBS for an hour at room temperature (RT), and incubated with primary antibody (Rabbit anti-PV, 1:1000, Swant, PV27a; rabbit anti-c-Fos, 1:500, Abcam, ab190289) overnight at 4℃. Then, sections were washed three times and incubated with secondary antibody (Jackson ImmunoResearch) and 4′,6-diamidino-2-phenylindole (DAPI, Santa Cruz Biotechnology, sc-3598) for two hours at RT in darkness. Finally, sections were washed and mounted (Antifade mounting medium, H-1000, Vector Laboratories). Images for virus and electrode location verification were acquired using an upright fluorescent microscope (Nikon Eclipse Ni-E). Images for $PV^+$ synapses quantification were acquired using confocal microscopy (Nikon A1HD25 high speed and large field of view confocal microscope).

##### Image analysis

$PV^+$ synapse expression (Fig 1) was quantified using FIJI/ImageJ software. The granule cell layer of DG, stratum pyramidale layer of CA1 and layer 2 of APC were set as regions of interest (ROIs) by visual inspection. A threshold value was determined by setting a gray value that separates background fluorescence from PV punctae and set for all images. 0.3-30 $\mu m^2$ was set as the size filter in the *analyze particle* tool. The percentage of $PV^+$ area and integrated density in each ROI were analyzed. Cell numbers (Supplementary Fig 1) were quantified by counting DAPI stained nuclei in the same ROIs. To smooth out the inhomogeneity of DAPI nuclear staining and separate the tightly packed nucleus, gaussian blur (sigma=3) and watershed segmentation were applied to preprocess images. Then, a threshold value was determined by setting a gray value that separates background fluorescence from gaussian blurred DAPI nuclei and set for all images. 20 μm$^2$ was set as the size filter in the *analyze particle* tool. The cell counts and area of each ROI were analyzed and used to calculate the cell density (cell density = counts / ROI area). For the DREADD virus specificity and function verification (Supplementary Fig 2), all mCherry-expressing neurons in the image were set as individual ROIs by using the *analyze particle* tool. A threshold was set for PV protein and c-fos protein separately. Then within each ROI, the co-labelling expression of mCherry and PV or mCherry and c-fos was checked by visual inspection.

### Supplementary Figures

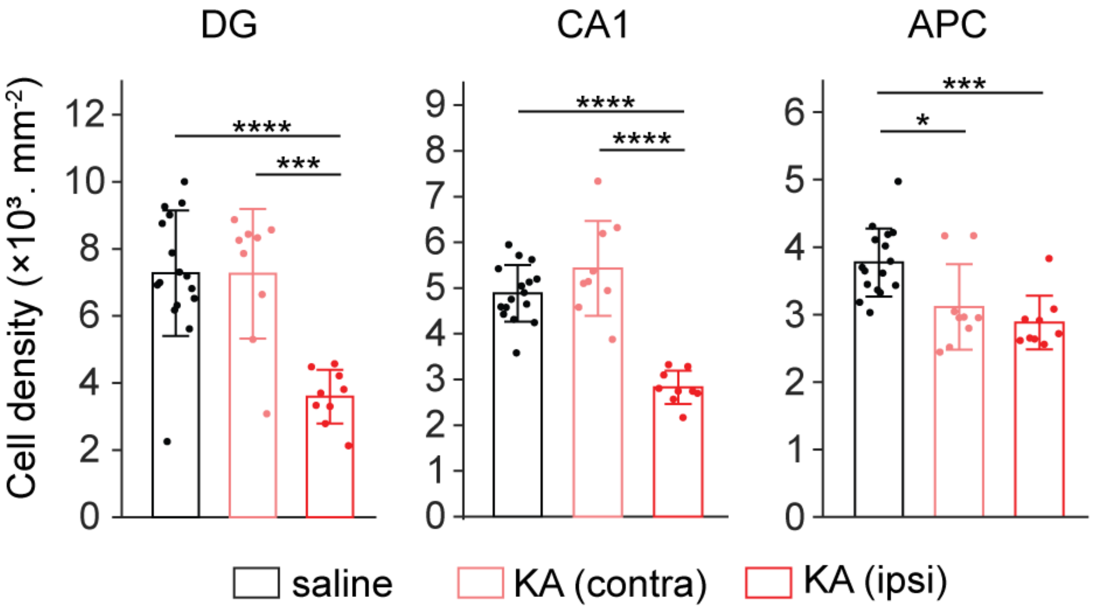


**Supplementary Figure 1. Neurodegeneration in HPC and APC in the chronic kainic acid model.** Quantification of cell density in DG (saline: 7274 ± 1869, n = 16; KA (contra): 7259 ± 1931, n = 9; KA (ipsi): 3592 ± 800, n = 9), CA1(saline: 4882 ± 618, n = 16; KA (contra): 5429 ± 1035, n = 9; KA (ipsi): 2826 ± 364, n = 9) and APC (saline: 3771 ± 503, n = 16; KA (contra): 3111 ± 633, n = 9; KA (ipsi): 2881 ± 396, n = 9). One-way ANOVA with Tukey multiple comparison as post hoc, * $p < 0.05$, *** $p < 0.001$, **** $p < 0.0001$. Data were shown as mean ± sd. n indicates the number of sections. N = 3 mice for each condition.

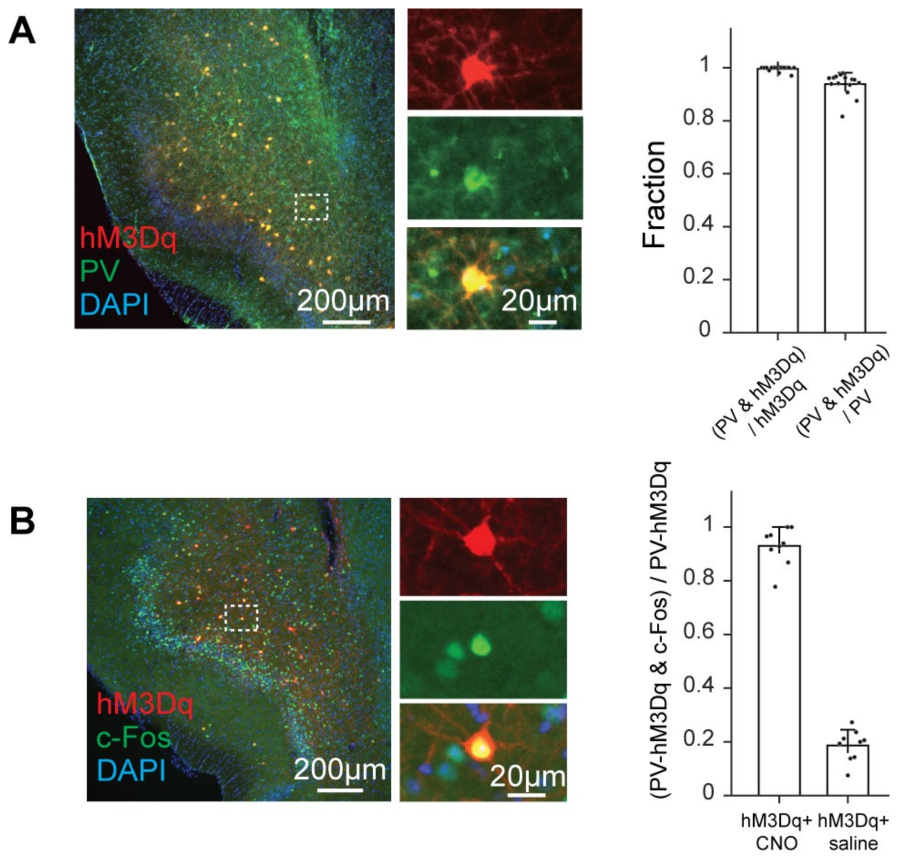


**Supplementary Figure 2. Specific expression and chemogenetic activation of PV interneurons by hM3Dq.** (**A**) Left, representative image showing hM3Dq-expressing neurons (red) and PV immunostaining (green) in the APC. The bar graph quantifies the fraction of hM3Dq-expressing neurons that are PV positive and the fraction of PV neurons expressing hM3Dq, demonstrating selective expression of PV interneurons. (**B**) Left, representative images showing c-Fos immunoreactivity (green) in hM3Dq-expressing PV interneurons (red). The bar graph quantifies the proportion of hM3Dq-positive PV interneurons that are c-Fos positive after CNO or saline administration, indicating robust activation of PV interneurons by chemogenetic stimulation.

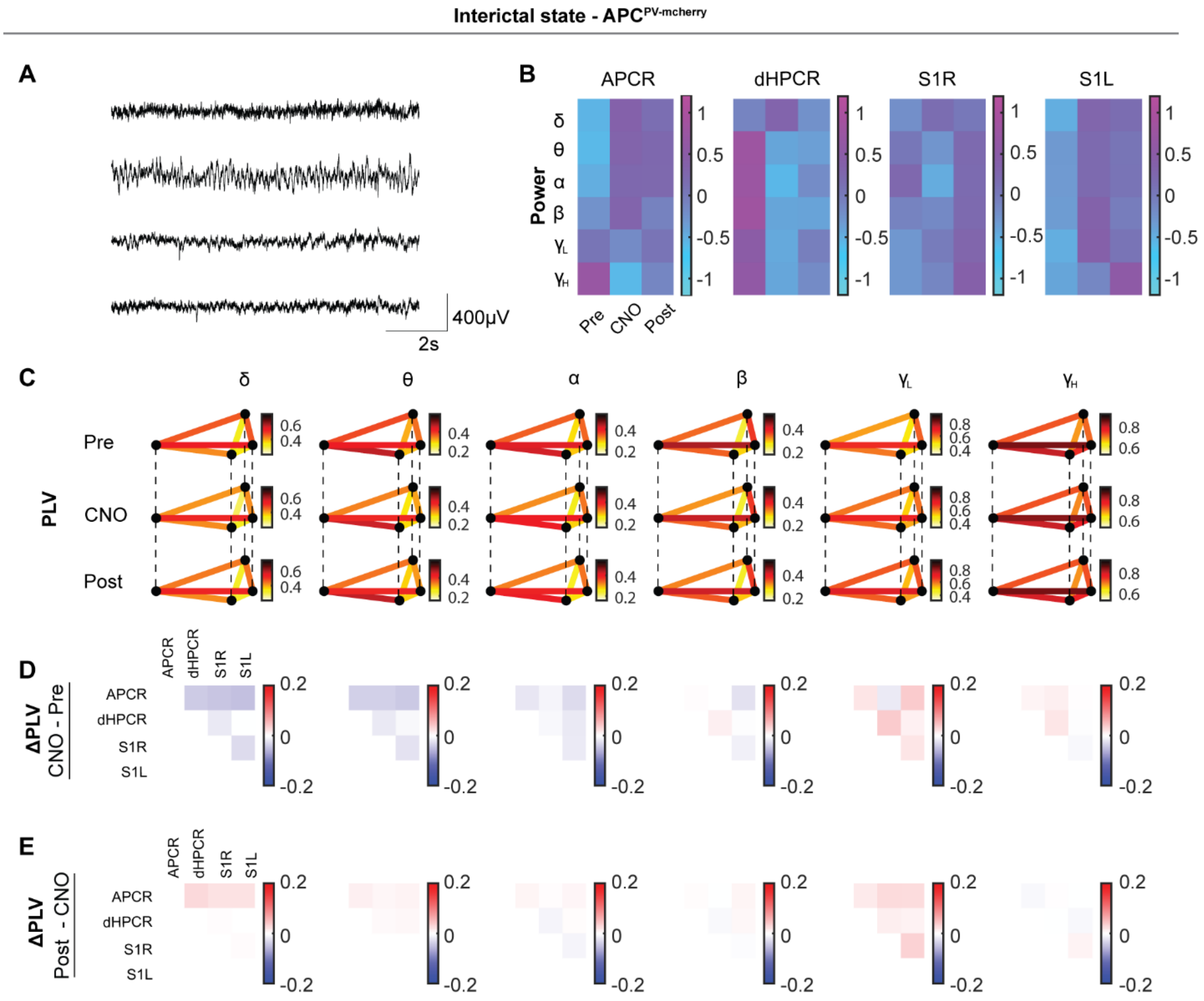


**Supplementary Figure 3. CNO administration on APC$^{PV\text{-}mcherry}$ mice fail to induce long-range network dynamics changes during interictal periods.** (**A**) Representative interictal LFP traces from four recording sites. (**B**) Normalized band power (z-scored within subjects for each channel and frequency band) during interictal periods displayed as a heatmap (rows: frequency bands; columns: Pre, CNO, and Post conditions; colors: mean band power; n = 7 mice). One-way repeated measures ANOVA. (**C**) Phase locking values (PLVs) between all region pairs across 6 frequency bands and Pre, CNO, and Post conditions (mean values, n = 7 mice). (**D, E**) ΔPLV matrices showing functional connectivity changes between: Pre and CNO conditions (D), CNO and Post conditions (E). One-way repeated measures ANOVA.

### Supplementary Table

**Supplementary Table 1. LFP features used in the PCA analysis**

| feature number | LFP feature name |
|---|---|
| 1 | power_APCR_delta_interictal |
| 2 | power_APCR_theta_interictal |
| 3 | power_APCR_alpha_interictal |
| 4 | power_APCR_beta_interictal |
| 5 | power_APCR_lo_gamma_interictal |
| 6 | power_APCR_hi_gamma_interictal |
| 7 | power_dHCR_delta_interictal |
| 8 | power_dHCR_theta_interictal |
| 9 | power_dHCR_alpha_interictal |
| 10 | power_dHCR_beta_interictal |
| 11 | power_dHCR_lo_gamma_interictal |
| 12 | power_dHCR_hi_gamma_interictal |
| 13 | power_S1R_delta_interictal |
| 14 | power_S1R_theta_interictal |
| 15 | power_S1R_alpha_interictal |
| 16 | power_S1R_beta_interictal |
| 17 | power_S1R_lo_gamma_interictal |
| 18 | power_S1R_hi_gamma_interictal |
| 19 | power_S1L_delta_interictal |
| 20 | power_S1L_theta_interictal |
| 21 | power_S1L_alpha_interictal |
| 22 | power_S1L_beta_interictal |
| 23 | power_S1L_lo_gamma_interictal |
| 24 | power_S1L_hi_gamma_interictal |
| 25 | plv_APCR-dHPCR_delta_interictal |
| 26 | plv_APCR-dHPCR_theta_interictal |
| 27 | plv_APCR-dHPCR_alpha_interictal |
| 28 | plv_APCR-dHPCR_beta_interictal |
| 29 | plv_APCR-dHPCR_lo_gamma_interictal |
| 30 | plv_APCR-dHPCR_hi_gamma_interictal |
| 31 | plv_APCR-S1R_delta_interictal |
| 32 | plv_APCR-S1R_theta_interictal |

| | |
|---|---|
| 33 | plv_APCR-S1R_alpha_interictal |
| 34 | plv_APCR-S1R_beta_interictal |
| 35 | plv_APCR-S1R_lo_gamma_interictal |
| 36 | plv_APCR-S1R_hi_gamma_interictal |
| 37 | plv_APCR-S1L_delta_interictal |
| 38 | plv_APCR-S1L_theta_interictal |
| 39 | plv_APCR-S1L_alpha_interictal |
| 40 | plv_APCR-S1L_beta_interictal |
| 41 | plv_APCR-S1L_lo_gamma_interictal |
| 42 | plv_APCR-S1L_hi_gamma_interictal |
| 43 | plv_dHPCR-S1R_delta_interictal |
| 44 | plv_dHPCR-S1R_theta_interictal |
| 45 | plv_dHPCR-S1R_alpha_interictal |
| 46 | plv_dHPCR-S1R_beta_interictal |
| 47 | plv_dHPCR-S1R_lo_gamma_interictal |
| 48 | plv_dHPCR-S1R_hi_gamma_interictal |
| 49 | plv_dHPCR-S1L_delta_interictal |
| 50 | plv_dHPCR-S1L_theta_interictal |
| 51 | plv_dHPCR-S1L_alpha_interictal |
| 52 | plv_dHPCR-S1L_beta_interictal |
| 53 | plv_dHPCR-S1L_lo_gamma_interictal |
| 54 | plv_dHPCR-S1L_hi_gamma_interictal |
| 55 | plv_S1R-S1L_delta_interictal |
| 56 | plv_S1R-S1L_theta_interictal |
| 57 | plv_S1R-S1L_alpha_interictal |
| 58 | plv_S1R-S1L_beta_interictal |
| 59 | plv_S1R-S1L_lo_gamma_interictal |
| 60 | plv_S1R-S1L_hi_gamma_interictal |
